# Seascape genetics at its finest: dispersal patchiness within a well-connected population

**DOI:** 10.1101/530451

**Authors:** C. Schunter, M. Pascual, N. Raventos, J. Garriga, J.C. Garza, F. Bartumeus, E. Macpherson

## Abstract

Dispersal is a main determining factor of population structure and variation. In the marine habitat, well-connected populations with large numbers of reproducing individuals are common but even so population structure can exist on a small-scale. Variation in dispersal between populations or over time is often associated to both environmental and genetic variation. Nonetheless, detecting structure and dispersal variation on a fine-scale within marine populations still remains a challenge. Here we propose and use a novel approach of combining a clustering model, early-life history trait information from fish otoliths, spatial coordinates and genetic markers to detect very fine-scale dispersal patterns. We collected 1573 individuals (946 adults and 627 juveniles) of the black-faced blenny across a small-scale (2km) coastline as well as at a larger-scale area (<50kms). A total of 178 single nucleotide polymorphism markers were used to evaluate relatedness patterns within this well-connected population. Local retention and/or dispersal varied across the 2km coastline with higher frequency of SHORT-range disperser adults; representing local recruitment; towards the southwest of the area. An inverse pattern was found for juveniles, showing an increase of SHORT-range dispersers towards the northeast. This reveals a complex but not full genetic mixing and suggests oceanic/coastal circulation as the main driver of this fine-scale chaotic genetic patchiness within this otherwise homogeneous population. When focusing on the patterns within one recruitment season, we found large differences in temperatures (from approx. 17oC to 25oC) as well as pelagic larval duration (PLD) for juveniles from the beginning of the season and the end of the season. We were able to detect fine-scale differences in HIGH-range juvenile dispersers, representing distant migrants, depending on whether they were born at the beginning of the season, hence, with a longer PLD, or at the end of the reproductive season. The ability to detect such fine-scale dispersal patchiness will aid in our understanding of the underlying mechanisms of population structuring and chaotic patchiness in a wide range of species even with high potential dispersal abilities.

## INTRODUCTION

In marine ecosystems, larval dispersal and recruitment are the main processes determining the population structure of species and understanding how they maintain connectivity is crucial for effective management (Jones et al. 2009). Although our understanding of larval dispersal in the marine environment has greatly improved over the last several decades (Cowen and Sponaugle 2009), much is still unknown. Most marine organisms exhibit a larval phase where small larvae can, in theory, travel large distances with ocean currents. Marine populations were thus generally thought to have elevated gene flow and little population structure (Mora and Sale 2002). This paradigm has been largely disproven as many recent studies have revealed that genetic population structure can exist across even small spatial scales and self-recruitment is often larger than previously believed (Carreras-Carbonell et al. 2007, Mokhtar-Jamai et al. 2011). In addition, direct measurements of dispersal using pedigree reconstruction analyses have revealed that self-recruitment and recruitment patterns are often heterogeneous and complex (Planes et al. 2009, Almany et al. 2017).

Dispersal patterns depend on pelagic larval duration (PLD) (Selkoe and Toonen 2011), reproductive output (Treml et al. 2012), ocean currents (White et al. 2010) or localized eddies and oceanographic retention patterns (Galarza et al. 2009, Schunter et al. 2011). The duration of the larval period is known to influence larval dispersal distance (Connor et al. 2007) and can have a direct relationship with connectivity (Treml et al. 2012, Pascual et al. 2017). Information on pelagic larval duration (PLD) has been used in numerous studies modelling dispersal capabilities, connectivity and for the establishment of networks of Marine Protected Areas (Mumby et al. 2011, Berumen et al. 2012, Melià et al. 2016, Almany et al. 2017). Several studies have shown an influence of ocean temperature on PLD (Connor et al. 2007), indicating that a change in temperature could have a direct influence on population connectivity (Kleypas et al. 2016). Considering the potential increase of sea temperatures due to global climate change (Marbà et al. 2015), studies analyzing the relationships among temperature, PLD and connectivity are essential in marine ecology.

In the last few years, studies of connectivity have used observations of kin, such as parent-offspring pairs or full siblings, to infer the trajectories and distances of larval movement at small scales, but require large sampling efforts and genotype datasets (Saenz-Agudelo et al. 2011, Harrison et al. 2012, D’Aloia et al. 2013, Schunter et al. 2014b, Melià et al. 2016, Almany et al. 2017). Further factors determining patterns of dispersal are availability of habitat, environmental selection, behavior or kinship (Gerlach et al. 2007). The population structure of marine organisms can be influenced by the degree to which larvae from different populations or demes are mixed in the plankton (Ben-Tzvi et al. 2012, D’Aloia and Neubert 2018). Some studies have demonstrated the existence of familial structure with marked genetic distances between recruitment cohorts; thus the lack of larval mixing during the planktonic period can lead to chaotic genetic patchiness (Iacchei et al. 2013, Bentley et al. 2014, Selwyn et al. 2016). Other studies, however, have shown that related individuals do not necessarily disperse cohesively, but limited dispersal was the cause of non-random patches of closely related individuals (D’Aloia et al. 2018). Furthermore, the effects of population structure and kinship on the evolution of social behaviors at sea have largely been ignored (Kamel and Grosberg 2013). Therefore, more research is needed in order to understand how dispersal, settlement and survival patterns of related individuals affect the structure of populations in the marine environment.

In this study, we analysed fine-scale structure and dispersal patterns of a rocky shore fish species, the black faced blenny (*Tripterygion delaisi*), using patterns of kinship inferred from single nucleotide polymorphism (SNP) genotypes. We also used a novel approach of binary data clustering to consider the effect of temperature on dispersal. Specifically, we combine larval traits, such as pelagic larval duration, and evaluate selective dispersal patterns via genetic marker heterogeneity in space. This approach is used to detect fine-scale patterns of dispersal of local and recent-migrant recruits, as well as differences between early and late season individuals in a well-connected population of *T. delaisi*, revealing patterns of relatedness that were not previous detected using traditional methods.

## METHODS

### Study system and sample collection

*T. delaisi* is a small, common, rocky-shore fish from the Mediterranean Sea and the eastern Atlantic (Carreras-Carbonell et al. 2006). Individuals of this species can live to a maximum of three years and adults display high territorial fidelity (Wirtz 1978, De Jonge and Videler 1989). Furthermore, 1-year old individuals are the most abundant component of the reproductive population. The spawning period starts in April, when water temperatures in the study area are ca. 15ºC and finishes at the end of July, with temperatures of ca. 23ºC (Raventós and Macpherson 2001). The planktonic larval duration of this species was estimated to be about 2-3 weeks, depending on the water temperature (Macpherson and Raventós 2006). Therefore, this species is a good study organism to analyse the relationships between temperature and dispersal patchiness.

Overall, 946 adult and 627 juvenile *T. delaisi* were collected on SCUBA from the coast of Blanes, Spain in the western Mediterranean Sea (41°40’30.7“N 2°48’14.4’E). Adults were caught with hand nets, body length was measured, and dorsal fin clips were taken non-lethally underwater, and then preserved in 95% ethanol. Adults were collected during the length of their breeding season from April 2010 to July 2010. Juveniles were also captured with hand nets on SCUBA between July and September 2010 and subsequently used for genetic and otolith analysis. Sea surface temperature readings were taken 5 m below the surface twice weekly with a Conductivity, Temperature and Depth probe (CTD) at Las Medes Marine Reserve (approximately 40km north of the focal sampling area in Blanes).

Two different spatial scales were studied. A ~2km stretch of coastline in the vicinity of Blanes (Figure 1) is the small-scale area in which exhaustive searches were performed for all dominant adult males protecting small nests (N=793), and randomized searches for non-dominant males and females (N=153), as well as juveniles (N=382). The large-scale area encompasses the coast towards the southwest and northeast of the small-scale area, spreading over a total of 42km, where juveniles (N=245) were collected (Figure 1). Juveniles were collected in this large-scale area from La Pilona (2.7922, 41.6701; N=1), Palomera (2.8078, 41.6781; N=62), Blanes north (2.8181, 41.686; N=13), Clotilde (2.8214, 41.6887; N=23), Lloret de Mar south (2.8402, 41.6929; N=50), Lloret de Mar north (2.86125, 41.699; N=23), Tossa de Mar (2.934, 41.7146; N=74). The exact location of collection of every sample in both areas was geo-referenced.

**Figure 1:**
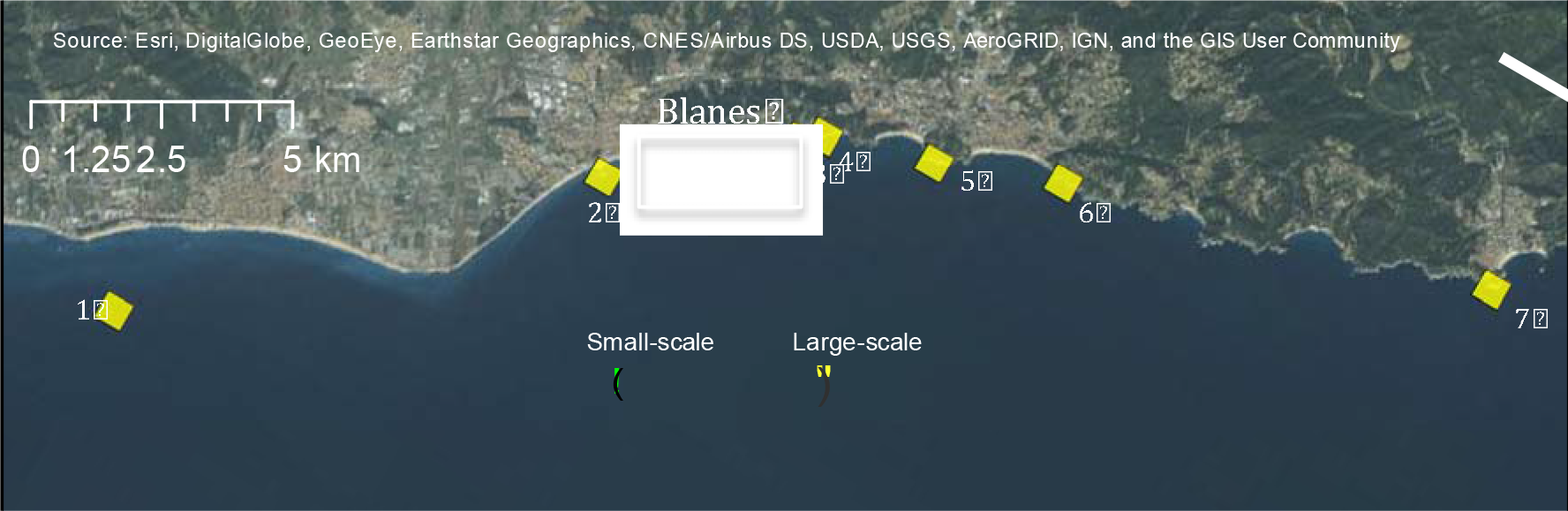
Map of sampling area including the small-scale intensive sampling area (Blanes) and the large-scale collection area. 1) La Pilona, 2) Palomera, 3) Blanes north 4) Clotilde, 5) Lloret de Mar south, 6) Lloret de Mar north, 7) Tossa de Mar.

### Otolith readings

Juvenile otoliths (lapilli) were extracted and mounted on microscope slides using thermoplastic glue (Crystalbond 509). To expose all daily increments, the otoliths were mounted and polished using two different grained sandpapers (3μm and 1μm Imperial lapping film, 3M) to obtain a thin section exposing the nucleus and the first growth ring. Reading of otolith bands was performed using a high-powered microscope with transmitted light (AxioPlan, Zeiss) connected to a ProgRes C10 camera (Jenoptic) and an image analysis system (Raventós and Macpherson 2005). To verify the first growth increment of the otolith, fertilized *T. delaisi* eggs collected in the field were reared in aquaria. First growth bands were checked at day 1 and day 3 post-hatching in ten individuals. This yielded both the timing of formation of the first band and the daily pattern of band deposition, and confirmed that band deposition took place daily from the day of hatching. Otolith marks in this species were always clear and belonged to Type Ia: an abrupt settlement mark with a sharp decrease in increment width across the settlement mark, completed within one increment (Wilson and McCormick 1999, Raventós and Macpherson 2001). The length of pelagic larval duration (PLD) for each individual was determined by counting the number of daily rings visible between the core and the settlement mark, and the age at sampling was determined by counting the total number of bands from core to margin. We analysed the otoliths along the longest radius from the centre to the edge and recorded otolith width during the pelagic larval period. We also recorded the size-at-hatching (radius from core to first increment) and size-at-settlement along the longest axis of the otolith. To minimize errors, all measurements were repeated three times. Trait differences were statistically evaluated in R (v. 3.3.1; RCoreTeam 2016).

A total of 200 randomly selected otoliths from juveniles collected in the small-scale area were read. Total body length of the juveniles correlated very well with the determined age by otolith reading (Pearson’s R = 0.9151, p < 0.001, Supplementary Material Figure S1), so the age of all 627 juveniles was estimated from body length with the regression equation. We inferred larval traits such as the hatching date for all juveniles by combining the age and the collection date of each individual. To compare the beginning and the end of the reproductive season, juveniles were assigned o two groups: ‘early juveniles’ if born between April and mid-May (N = 164 within Blanes and N = 224 for the large-scale area) and ‘late juveniles’, born between the 15^th^ of June and the end of the recruitment season, which is at the beginning of September (N= 121 in Blanes and N= 229 for the large-scale area). Inferred larval traits (size at hatching, size at settlement and pelagic larval duration) were compared between early juveniles (N=78) and late juveniles (N=41) with otolith readings by t-tests in R.

### Genetic analyses

All 1573 individuals were genotyped with 192 SNP markers developed for *T. delaisi*; details of the SNP development can be found in (Schunter et al. 2014b). Markers that were not in Hardy-Weinberg or linkage equilibrium were discarded and the remaining 178 SNP loci used in subsequent population genetic and parentage analyses, as described in Schunter et al. (2014b). Heterozygosities for each marker and all markers combined were computed with GenAlEx (v. 6.5; Peakall and Smouse 2012). Pairwise relatedness values (r-values) were obtained by using the Queller and Goodnight equation (Goodnight 1989) implemented in the SPAGeDi software (Hardy and Vekemans 2002). To find patterns of high or low relatedness amongst thousands of pairwise relatedness values, we used a relatedness ‘ratio’ to identify individuals with large numbers of high relatedness values. For each individual, the number of r-values above 0.1 was counted and divided by the total number of comparisons. The top 25% of individuals with the highest ratios were termed ‘locals’, whereas the samples with the lowest ratios (bottom 25%) were referred to as ‘recent migrants’. Differences in spatial structure of these groups of individuals were evaluated using their geo-referenced locations of collection and with a Kolmogorov-Smirnov test in R.

### Spatial and Genetic Clustering

To evaluate the spatial distribution of kinship we incorporated spatial structuring of samples into the analysis. Latitudes and longitudes for each individual’s sampling location were used to find spatial clusters within the small-scale (Blanes) area as well as in the large-scale sampling area. We found a subset of spatial clusters, with the highest likelihood score, with the R package EMCluster (Chen & Maitra, 2015) and assigned individual geo-localized samples to the respective spatial clusters.

To take into account the spatial signature of relatedness estimates, we averaged individual genetic distances considering their spatial clustering across the study area. Specifically, we computed the genetic distances of each individual with each of its conspecifics in the same spatial cluster and with all other individuals (see Figure 2). As pair-wise relatedness values can be negative when using an unbiased estimator, especially in continuous large populations (Amos et al. 2001) we first converted each single relatedness value into a distance between 0 and 1 with the formula (d=1−(r−a)/(b−a)), where a and b are the minimum and maximum value of relatedness (i.e., r), respectively. Thus, for each individual we computed two distinct values of Mean Genetic Distances (MGD): (i) the mean genetic distance of an individual in a given spatial cluster to all the other members of the same cluster (MGDin), and (ii) the mean genetic distance of that individual to the rest of individuals outside of its spatial cluster (MGDout). Finally, we characterized each individual with these two measures and performed an Expectation-Maximization binary clustering with the R package EmBC (Garriga et al. 2016). This analysis allowed us to partition all individuals according to four categories (Figure 2): 1. MIXED-range dispersers: individuals with low MGD with respect to individuals in their own spatial cluster (MGDin) and low MGD with respect to individuals in other spatial clusters (MGDout), which are most likely individuals with relatively common genotypes that confer medium relatedness values with many other individuals. 2. SHORT-range dispersers: individuals with low MGDin and high MGDout, which therefore are the individuals that have high relatedness to neighboring individuals. 3. MEDIUM-range dispersers: individuals with high MGDin and low MGDout, which are fish that came from the area but from a different spatial cluster. 4. HIGH-range dispersers: individuals with high MGDin and high MGDout, which means that these individuals have very low relatedness with most of the other samples and could be long-distance dispersers from a different genetic population.

**Figure 2:**
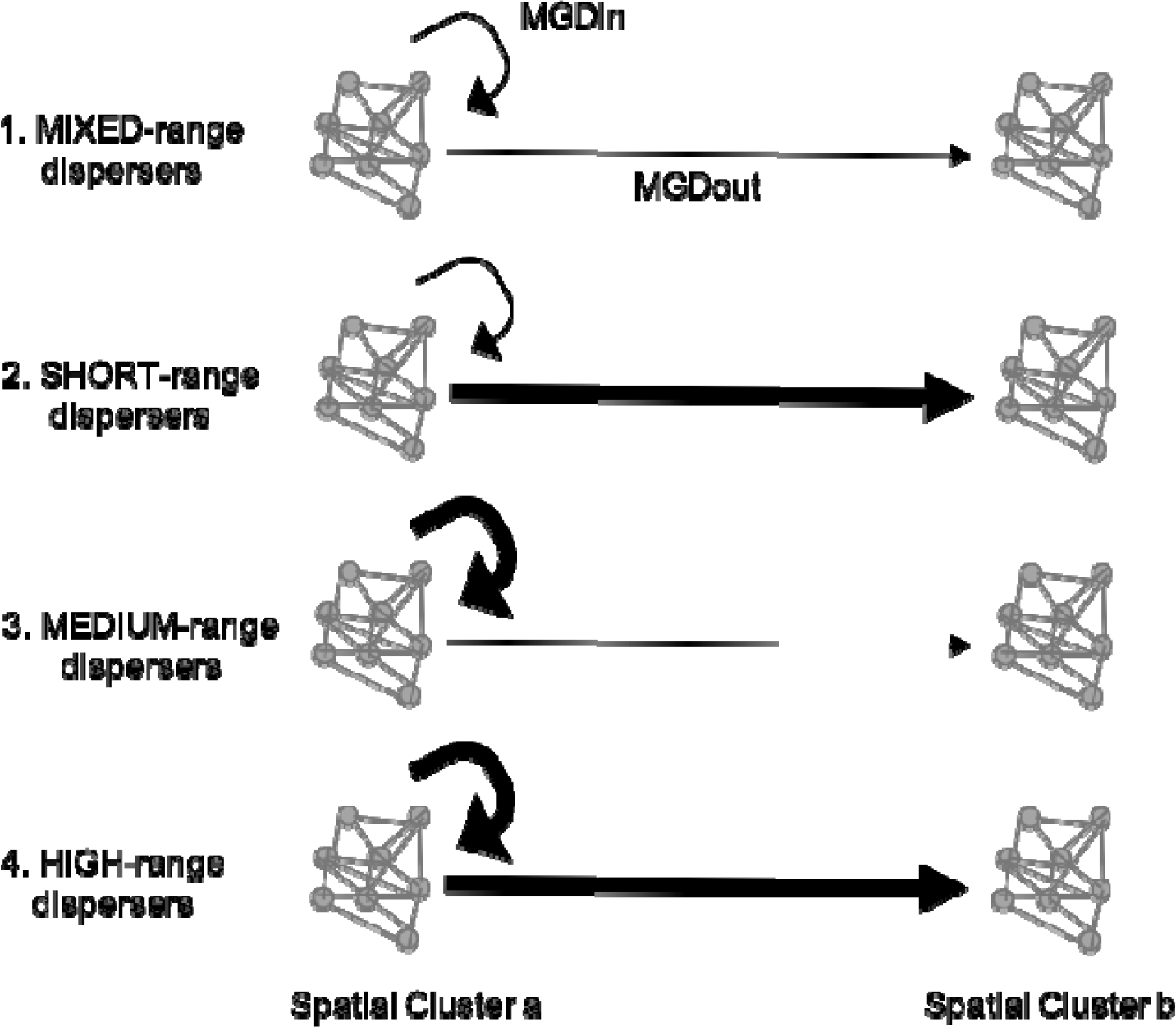
Conceptual Plot showing the four categories of dispersers. Two spatial clusters are represented and the mean genetic distance (MGD) among the individuals within one cluster (MGDin) indicated with a circular arrow and with individuals of other spatial clusters (MGD out) with the straight arrow. The thickness of the lines represents the distance: thick= high MGD, thin=low MGD.

## RESULTS

### Genetic and phenotypic diversity

We determined the hatch date of each individual and the number of days the larvae spent in the water column with the subset of 200 randomly selected otoliths. The amount of time a larva spent in the water column depended on when in the reproductive season it was born (Pearson’s r = −0.6057, p < 0.001; Figure S2). Individuals hatching in April can have a planktonic tenure of up to 10 days longer than those hatching in July. No significant differences were found between ‘early juveniles’ and ‘late juveniles’ for size-at-hatching (t = 1.73, p = 0.086) or size-at-settlement (t =−1.009, p = 0.32). Average PLD was 19.5 days for early juveniles and 16 days for late juveniles and were significantly different (t = 9.19, p = 2.596e−14). The data were found to be normally distributed and no significant differences were found between recent migrant and local juveniles, as categorized based on the relatedness ratio nor for any otolith characteristic (size-at-hatching t = 0.57, p = 0.56; size-at-settlement t = 0.25, p = 0.80; PLD t = −0.08, p = 0.94).

No significant differences in heterozygosity were found between juveniles and adults (t = −0.53, p = 0.59) nor between early and late juveniles (t = −0.95, p = 0.34) for all loci combined (Figure S3). Small differences in heterozygosity between early and late juveniles were found when analyzed locus by locus (Figure S4). Relatedness also did not differ between adults and juveniles (t = 1.301, p-value = 0.194; Figure S5). Due to a longer PLD at the beginning of the season, individuals could potentially travel further distances and disperse further and we might expect differences in the genetic distance/relatedness of these individuals, given that *T. delaisi* larvae are able to travel 11.5km in 10 days (Schunter et al. 2014b). However, the average relatedness values of early juveniles (−0.00099±0.015) and late juveniles (−0.00027±0.015) were not significantly different (t=−0.55, p-value=0.58) nor did the relatedness value distribution differ between adults and juveniles (Figure S6). Of the 157 juvenile individuals categorized as ‘locals’ due to high relatedness ratios, 29 were early juveniles and 17 late juveniles, whereas of the 157 juvenile individuals considered ‘recent migrants’, with low relatedness ratios, 19 were early and 17 late juveniles. Interestingly, one of the individuals considered a ‘local’ is the sole sample from la Pilona (Figure 1) in the southernmost part of the large-scale sampling area.

We did not find differences in spatial distribution patterns between adults and juveniles representing two different years of recruitment (Kolmogorov-Smirnov D = 0.438, p = 0.09). Immigration of less-related individuals, possibly from more distant populations, into the small-scale area could potentially result in different spatial patterns for these ‘recent migrants’. Adult migrants were differently distributed when compared to the rest of adults (D = 0.72, p < 0.001) with some differences evident in the southernmost part of the distribution (Figure S7). However, ‘recent-migrant’ juveniles were not distributed differently from the rest of the juveniles (D = 0.25, p = 0.6994), nor were spatial differences found between early and late ‘recent-migrant’ juveniles in the small-scale area (D = 0.125, p = 0.996; Figure S8).

### Fine-scale spatial clustering analysis

With hundreds of pairwise relatedness comparisons per individual, it is difficult to detect fine-scale genetic structure, although there could be spatial clusters. To evaluate genetic clustering in space, we first found significant spatial clusters in the small-scale area for both adults and juveniles (Figure 3). We found six spatial clusters for the adults and seven in the juvenile samples. The additional cluster found for juveniles (SplClust6; Figure 3b) is in an area that could correspond to non-suitable habitat for *T. delaisi*, as we found no adults and only a few juveniles.

**Figure 3:**
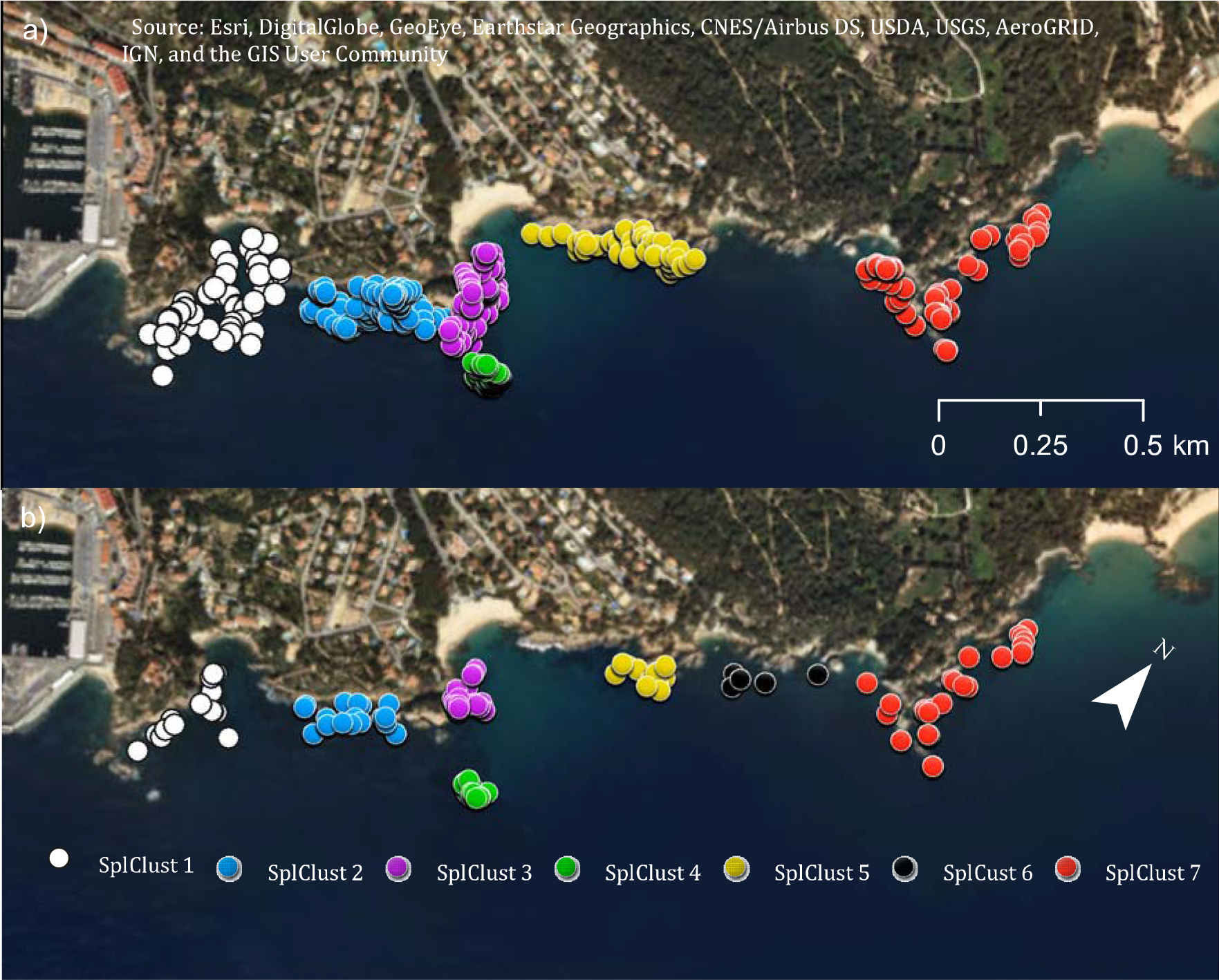
Spatial clusters within the small-scale area for a) adults and b) juveniles. Spatial cluster 6 only exists for the juveniles and few individuals were found.

The proportion of individuals in each cluster that came from the SHORT-range disperser category varied from southwest to northeast along the coastline (Table S1). For juveniles, the frequency of SHORT-range dispersers increased along the coast from the southwest to the northeast, where they constituted 81% of all fish sampled, whereas the pattern was reversed for adults; that is, the frequency of SHORT-range dispersers decreased from southwest to northeast (Figure 4). In contrast, HIGH-range dispersers, which could be considered immigrants, were less frequent than SHORT-range dispersers at both life stages/years of recruitment (adults likely recruited in 2009 and juveniles in 2010) and were relatively homogeneously distributed along the coast (Figure 4).

**Figure 4:**
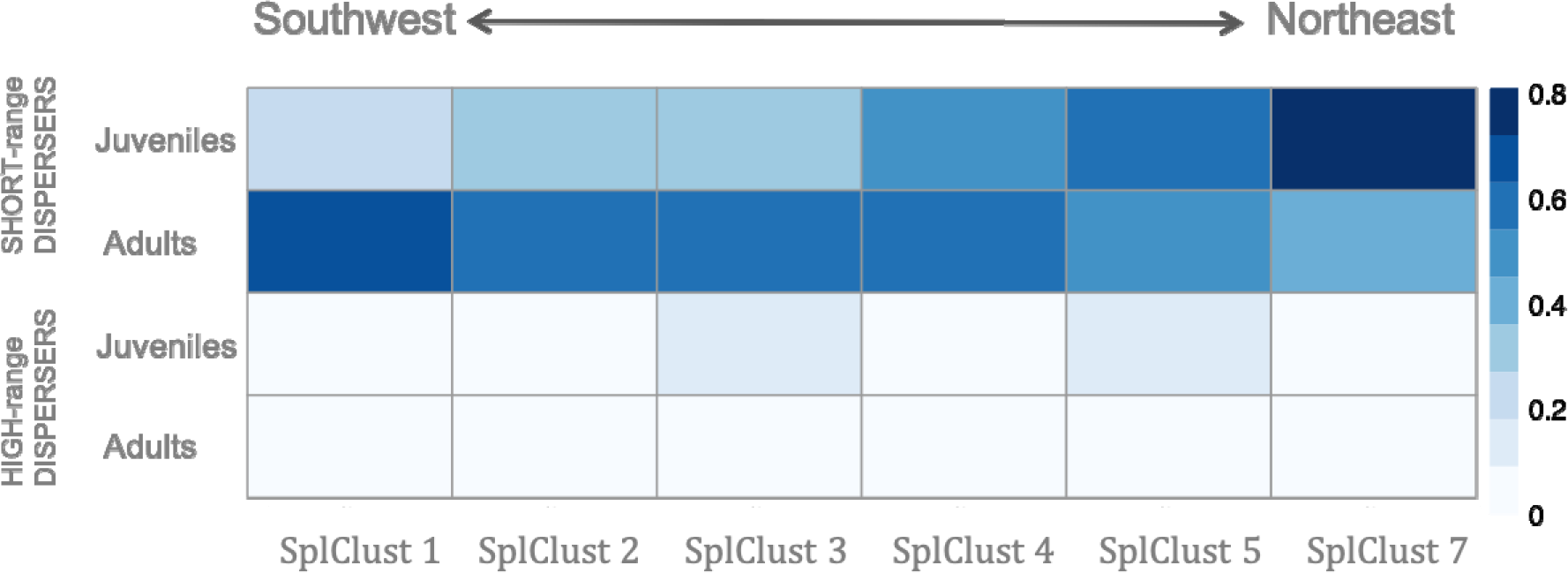
Proportion of adults and juveniles in each spatial cluster (SplClust; Figure 3) in the SHORT-range disperser category, which are individuals with low mean genetic distance (MGD) with respect to their own spatial cluster and a high MGD with respect to other spatial clusters as well as HIGH-range dispersers, which have high MGD to all clusters. SplClust6 is not included because it was only found in juveniles and in very low frequency.

Juveniles born at the beginning of the season (early juveniles) and at the end of the season (late juveniles) have differences in pelagic larval duration (Fig S2), and different patterns were also observed for specific spatial clusters (Table 1). Spatial Cluster 4 (Figure 3) is a rocky reef patch called ‘La Roca’ that is separated from the coastline by sand and has the furthest distance from shore. In Spatial Cluster 4, at the beginning of the season (with longer PLDs), only HIGH-range disperser juveniles were found (Figure 5). These individuals could be immigrants dispersing from distant populations. Late juveniles, with shorter PLDs, on the other hand, had a higher proportion of SHORT-range dispersers compared to HIGH-range dispersers at La Roca (SplClust4) indicating a more local origin. Most HIGH-range dispersers in late juveniles were found in spatial clusters 2 and 5 (Figure 5).

**Figure 5:**
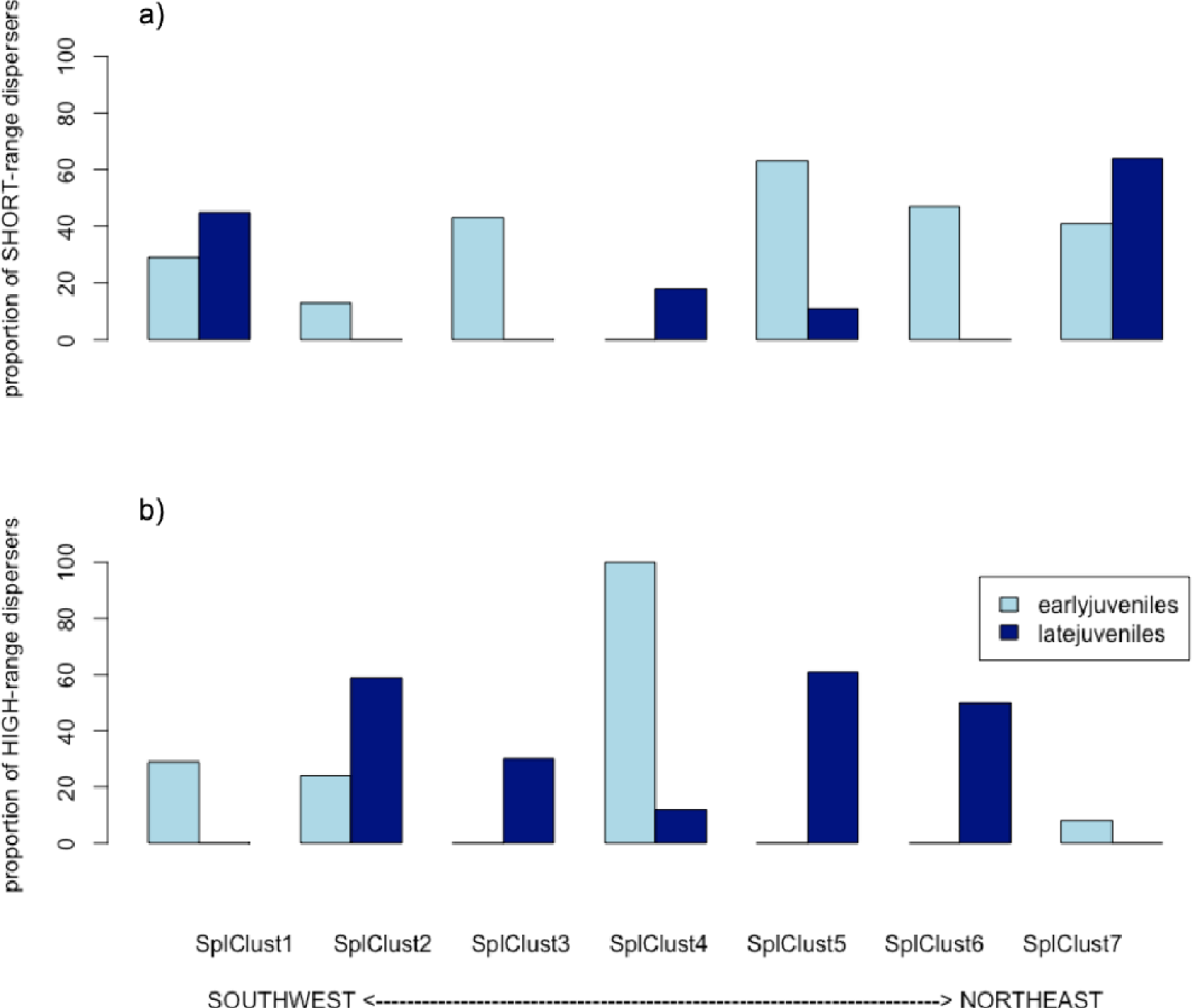
Proportion of low dispersers (a) which are individuals with a low mean genetic distance (MGD) with respect to their own spatial cluster and a high MGD with respect to other spatial clusters and high dispersers (b) which are individuals with high MGD with respect to all clusters, in early and late juveniles in each spatial cluster (from Figure 3).

**Table 1:** Proportion of individuals that belong to each spatial cluster in the large-scale sampling area. The clusters correspond to Figure 6.

| **Juveniles large scale area** | SplClust1 | SplClust 2 | SplClust 3 | SplClust 4 | SplClust 5 |
| --- | --- | --- | --- | --- | --- |
| N | 98 | 119 | 174 | 79 | 157 |
| MIXED-range dispersers | 0 | **0.62** | 0.3 | 0.07 | 0.03 |
| SHORT-range dispersers | **0.7** | 0.34 | **0.55** | **0.75** | 0.1 |
| MEDIUM-range dispersers | 0.3 | 0.04 | 0.16 | 0.08 | **0.84** |
| HIGH-range dispersers | 0 | 0 | 0 | 0.1 | 0.03 |

*MIXED-range dispersers: individuals with low Mean Genetic Distance (MGD) with respect to individuals in their own spatial cluster (MGD<sub>in</sub>) and low MGD with respect to individuals in other spatial clusters (MGD<sub>out</sub>). SHORT-range dispersers: individuals with low MGD<sub>in</sub> and high MGD<sub>out</sub>. MEDIUM-range dispersers: individuals with high MGD<sub>in</sub> and low MGD<sub>out</sub>. HIGH-range dispersers: individuals with high MGD<sub>in</sub> and high MGD<sub>out</sub>. Values above 0.5 are in bold.*

### Large-scale spatial clustering analysis

Genetic spatial clustering for juveniles across the whole large-scale sampling area, spanning a distance of almost 50km, identified five distinct clusters, four of them including individuals within the small-scale area (Figure 6). Some of the clusters of the large-scale analysis combined clusters identified in the small-scale area. For instance, SplClust 2 of the large-scale analysis (Figure 6) included all individuals of SplClust 2, 3 and 4 in the small-scale analysis (Figure 3).

**Figure 6:**
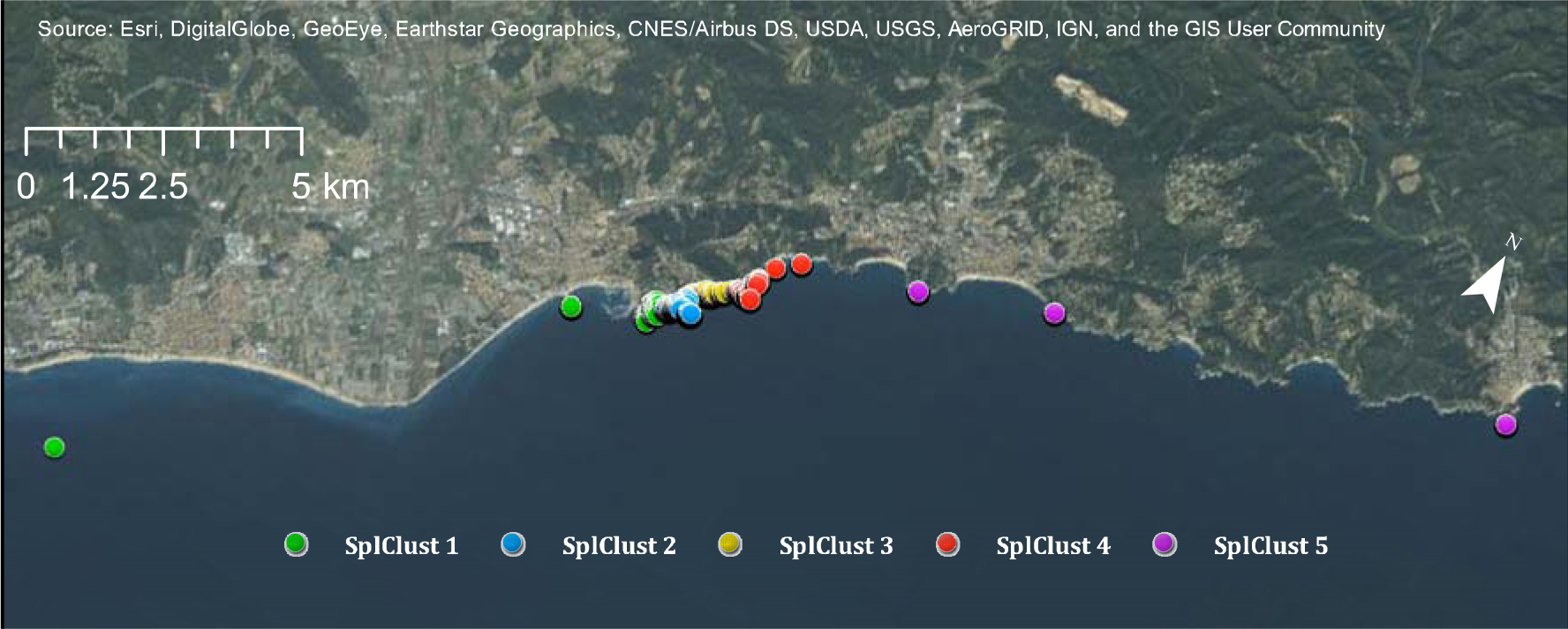
Spatial clusters for juveniles within the whole large-scale area

The proportion of HIGH-range dispersers, and thus likely immigrants, was very low for the large-scale dispersal area (Table 1). As the large-scale area is almost 50km of coastline, these individuals would have to come from even more distant locations. All clusters had a higher proportion of SHORT-range dispersers than of MEDIUM or HIGH-range dispersers, except Spatial Cluster 5, which is the northeastern-most sample area, which had a higher proportion of MEDIUM-range dispersers (Table 1), meaning that individuals in this cluster had less genetic relatedness to each other and were more related to the other locations from the southwest.

## DISCUSSION

### Dispersal and recruitment patterns in the black-faced blenny population

In the present work, we document very fine-scale patterns of dispersal and recruitment in a population of the black-faced blenny (*Tripterygion delaisi*) on the northern Mediterranean coast of Spain, through the combined use of genetic relatedness indices and a novel approach of binary data clustering that incorporates spatial patterns. In such a well-connected population with large numbers of dispersing individuals, genetic homogeneity would be expected and no population structure or population-wide pairwise relatedness was found with traditional methods (Schunter et al. 2014b). However, binary clustering of genetic relatedness indices mapped into space revealed a slight clustering of *T. delaisi* individuals with different dispersal patterns. More adult blennies (sampled in 2010 and recruited mostly in 2009) considered short distance dispersers, were found in the southwestern parts of the small-scale study area. In contrast, juvenile individuals that recruited and were sampled in the following year (2010), had more short distance dispersers in the northeastern part of the 2km coastline. Previous population genetic studies of this species with microsatellite loci, in the same small-scale area and the surrounding areas (up to 50km), identified only one genetic unit with high within-population self-recruitment (Carreras-Carbonell et al. 2006, 2007). Sibship analysis of *T. delaisi* from the same location found that full siblings can recruit many km apart from each other, further indicating a genetically homogenous population (Schunter et al. 2014b). Nonetheless, the binary clustering analysis across the large-scale area demonstrated that a large proportion of juveniles collected northeast of the small-scale sampling region were MEDIUM-range dispersers and thus more related to the juveniles of this sampling region. This could potentially be the result of a previously unknown directional movement of larvae from southwest to northeast. The detection of fine-scale clusters at both small and large-scale study areas potentially associated with oceanic current patterns shows that there is a complex but not full genetic mixing and suggests oceanic/coastal circulation as the main driver of the fine-scale chaotic genetic patchiness within this otherwise homogeneous population.

### Larval dispersal by oceanic/coastal current drives fine-scale chaotic patchiness

Chaotic genetic patchiness, a term coined decades ago (Johnson and Black 1982), has become commonly used to describe patterns of genetic diversity and population structure (Selkoe et al. 2006, Iacchei et al. 2013, Bentley et al. 2014). Broquet *et al.* (2013) conclude through simulations that chaotic genetic patchiness could occur by a combination of genetic drift due to small effective size and collective dispersal at the larval stage. Such collective dispersal would most likely occur through retention of local larvae near the natal site and close to shore, as suggested by hydrodynamic modelling of larval dispersal (James et al. 2002). Even though chaotic genetic patchiness is generally associated with marked genetic distances between cohorts our results seem to be consistent with the dispersal hypothesis as the main driver of our fine-scale genetic patchiness. In essence, the higher degree of relatedness in the southwestern portion of the study area could be a result of a physical retention process of ‘locals’ at settlement or a collective movement after settlement. Importantly, these aggregations in space vary across two recruitment seasons, an additional sign of dispersal-driven fine-scale chaotic genetic patchiness.

The reproductive biology of *Tripterygion delaisi* could suggest that other mechanisms may play a minor role in generating fine-scale patterns of chaotic genetic patchiness. For this species, the number of adults (i.e., potentially reproductively active) is very large, with more than 1000 individuals on a narrow 2km stretch of coastline (Schunter et al. 2014b). Reproductively active individuals include females and two types of males, the dominant nest-making male and the sneaker male (De Jonge and Videler 1989). So-called sweepstake reproduction has been shown to cause genetic patchiness in other species (Iacchei et al. 2013, Pusack et al. 2014) and for *T.delaisi* it could be that a few successful (i.e. the dominant) males contribute to the majority of offspring. However, the testes of *T. delaisi* sneaker males are significantly larger than those of dominant males and gene expression analysis found genes involved in differentiation and development, suggesting that these males are reproductively active (Schunter et al. 2014c). Moreover, in our analyses, we did not find any evidence of sweepstakes reproductive success, as genetic diversity did not decrease between adults and juveniles within the breeding season.

Furthermore, relatedness did not differ for juveniles recruiting at the beginning or at the end of the breeding season. Sweepstakes-driven patchiness thus seems unlikely, but cannot be ruled out, as high variability in the number of contributing adults over time has been shown in other marine fishes (Pusack et al. 2014, Waples 2016).

Patchiness could also be due to recruitment differences related to the conditions encountered by larvae. Pelagic larval duration (PLD) varied significantly with the hatching date of the larvae, with PLD up to 10 days shorter at the end of the season than at the beginning. Larvae that spend more time in the water column have more opportunities to disperse further and might include more immigrants from other populations. Yet, a similar proportion of *T. delaisi* migrants arrived early in comparison to late in the season in the whole sampling area. Nonetheless, at ‘La Roca’ (Spatial Cluster 4), a submerged rocky habitat not continuous with the rocky shoreline habitat, all early juveniles collected (N=29) were HIGH-range dispersers and can be considered migrants. This could hint at collective dispersal, at least at the beginning of the season, which Broquet and coauthors (2013) suggested as the second source for chaotic genetic patchiness and greater than expected levels of kinship as indicator have been found in other marine organisms (see Eldon et al. 2016). However, a previous study only found one full sibling pair of *T. delaisi* settling together in the same area (Schunter et al. 2014b) and the fine-scale patchiness pattern might be due largely to seasonal variation in current patterns.

Increased ocean temperatures, due to global climate change, have been suggested to decrease connectivity between populations and increase self-recruitment due to a general decline in pelagic larval duration (PLD) at higher temperature (Green and Fisher 2004, Connor et al. 2007, Munday et al. 2009, Kendall et al. 2016). The effect of temperature on larval connectivity has also been modelled and results suggested a considerable change in larval recruitment and connectivity (Lacroix et al. 2017). Such changes in dispersal potential and connectivity in *T. delaisi* could have detrimental effects on population genetic diversity. In our study area, the difference in sea surface temperature between the beginning (April) and the end of the reproductive period (July) is approximately 8º C.

This is larger than the predicted temperature increase in the western Mediterranean Sea (ca. 0.25ºC per decade) towards the end of the present century due to climate change (e.g. Marbà *et al.* 2015). Mean PLD of the black-faced blenny clearly decreases with increased temperature, but in the small-scale area studied here, this has little effect on the overall connectivity of the population since the proportion of individuals arriving from distant localities is similar regardless of PLD or water temperature. Even with a possible reduction in PLD over time due to climate change, intra-annual variation in conditions encountered by larval *T. delaisi* may provide a buffer and allow maintenance of generally well-connected populations.

### Conclusion

The species *Tripterygion delaisi* on the Mediterranean coast of Spain is a well-connected unit but appears to have fine-scale structure in both space and time. The combination of geo-localized individual locations, hatching date information, and genotypic SNP data allowed detection of patches of larvae with particular dispersal histories. Yet, very fine-scale genetic patchiness across space and time was only apparent when using binary clustering techniques, previously used primarily for animal movement trajectory segmentation (Garriga et al. 2016). The ability to detect such fine-scale dispersal patchiness will aid in our understanding of the underlying mechanisms of population structuring and chaotic patchiness in a wide range of species, including those with high potential dispersal abilities.

## ACKNOWLEDGEMENTS

Collection and field procedures followed the Spanish Laws (Royal Executive Order, 53/2013) for Animal Experimentation, in accordance with the European Union directive (2010/63/UE).

SNP assay sequences are deposited at the NCBI dbSNP with accession numbers 778235193 to 778235848. Genotypes of *Tripterygion delaisi* adults and recruits of the 178 SNP markers can be found in the Supplementary Materials Table S2. R scripts of the EMbC analyses can be found as Supplementary Data 1.

We are grateful to the Southwest Fisheries Science Center Molecular Ecology and Genetic Analysis Team for support in the lab and helpful comments and Jinliang Wang for his help in converting relatedness into a distance measurement. This work was partially funded by the Spanish Ministry of Science and Innovation through the CTM2017-88080 (AEI/FEDER, UE) project. The authors CS, MP and EM are part of the research groups 2017SGR-1120 and 2017SGR-378 of the Generalitat de Catalunya and CS was funded at the time of data collection by a JAE-Predoctoral Fellowship.

