## Supplementary Data for "Seascape genetics at its finest: dispersal patchiness within a well-connected population"

Supplementary Materials

Figure S1:

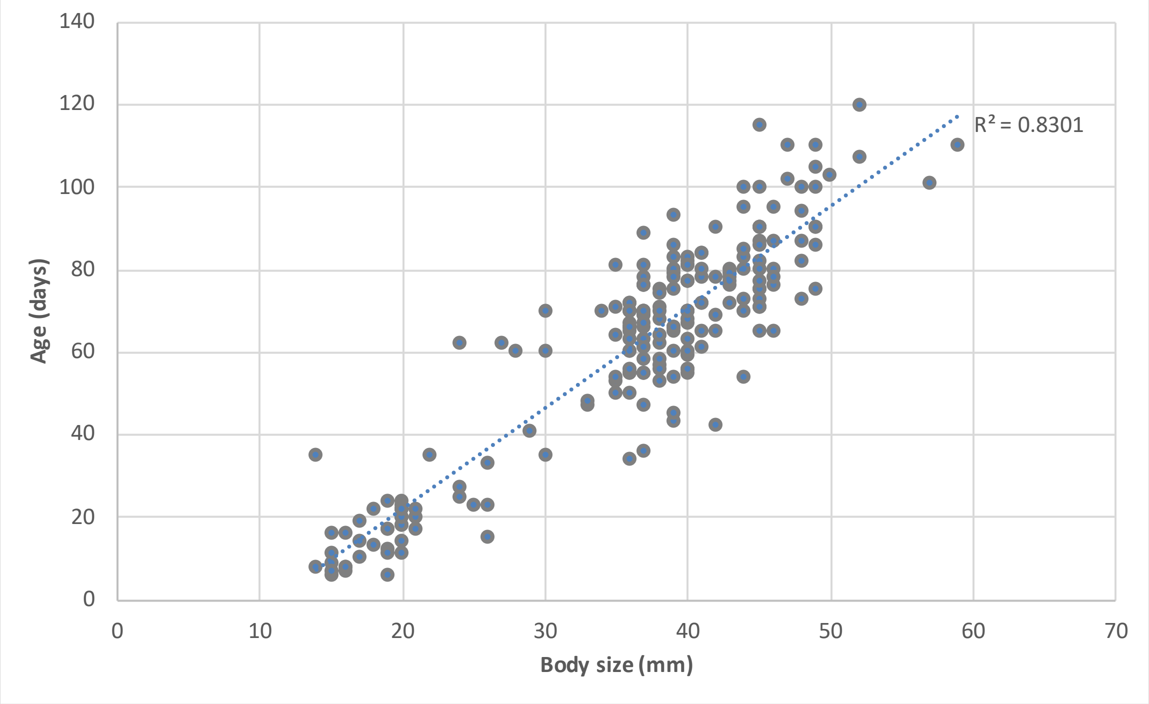


**Figure S1**: Correlation between body size and age according to otoliths reading for the 200 randomly selected juveniles.

Figure S2:

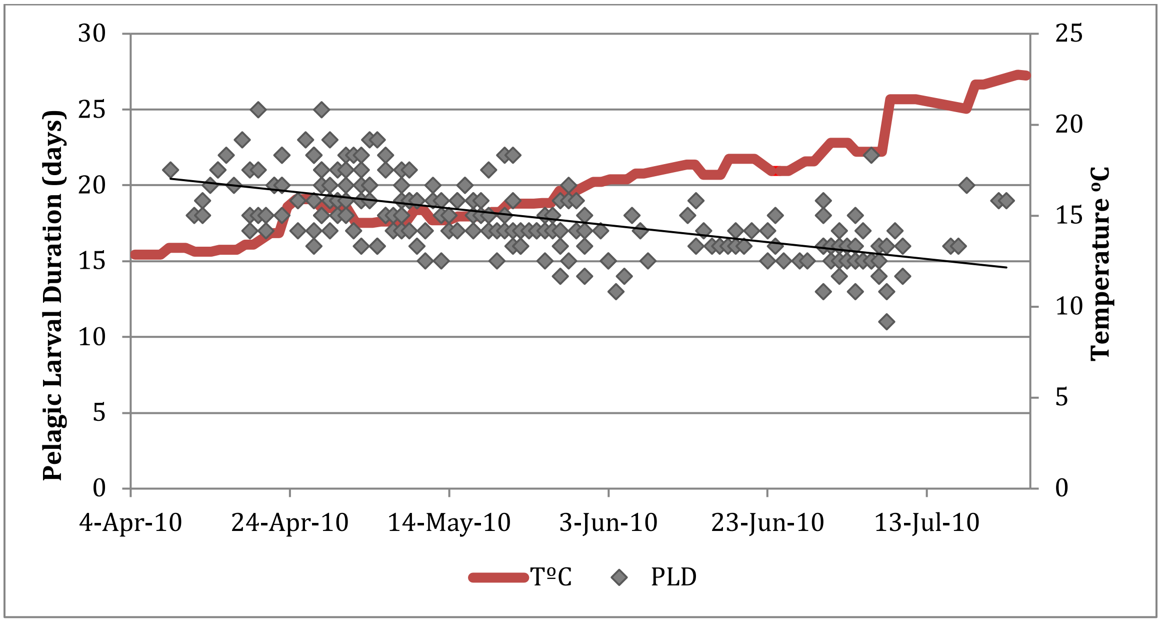


**Figure S2***:* Correlation between pelagic larval duration (PLD) and the date of juvenile hatching established by otolith readings for the 200 randomly selected juveniles. Sea temperature at 3 meters depth increases across time during the studied area. The significance of the correlation was Pearson's r= -0.6057, p<0.001.

Figure S3:

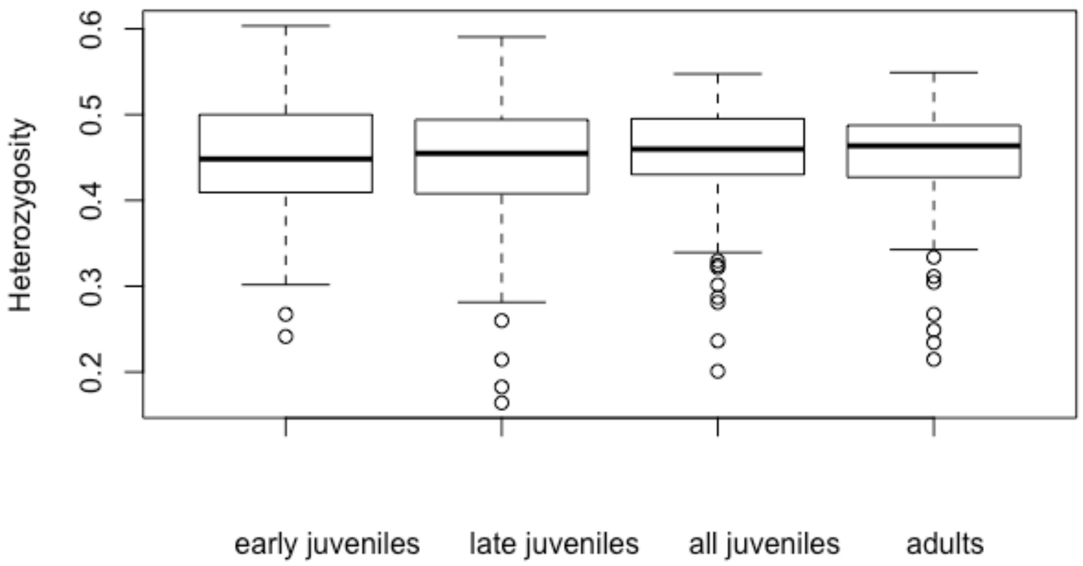


**Figure S3**: Heterozygosity boxplots for different life stage groupings. No difference in heterzygosity shows that there is no evidence of any sweep stake event.

Figure S4:

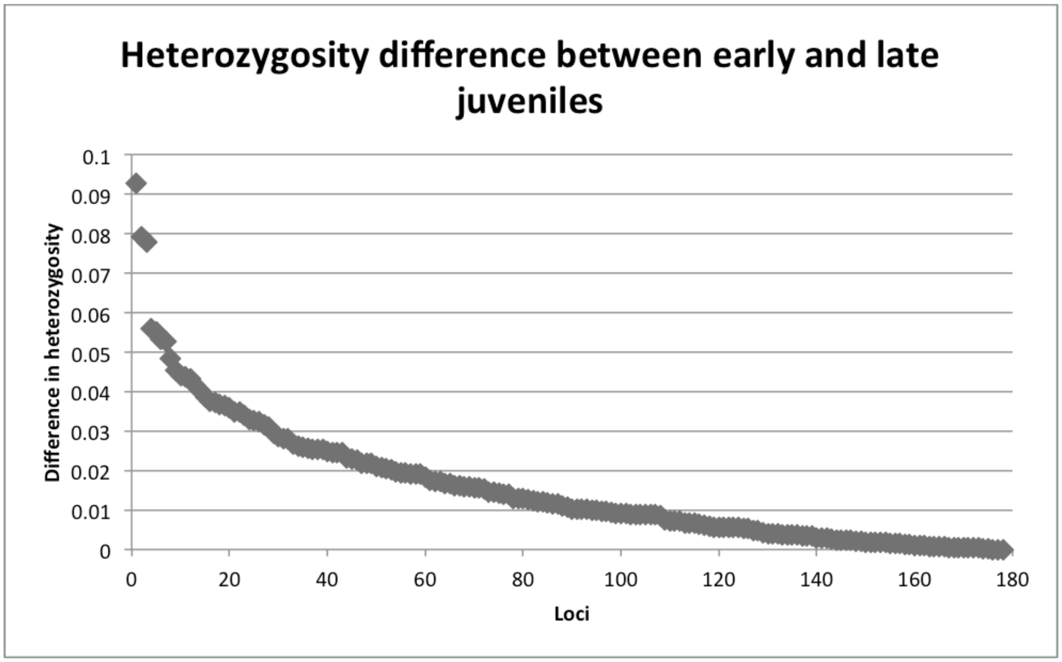


**Figure S4**: Absolute differences in heterozygosity between early and late juveniles for each of the 178 loci ordered from more to less differentiated.

Figure S5:

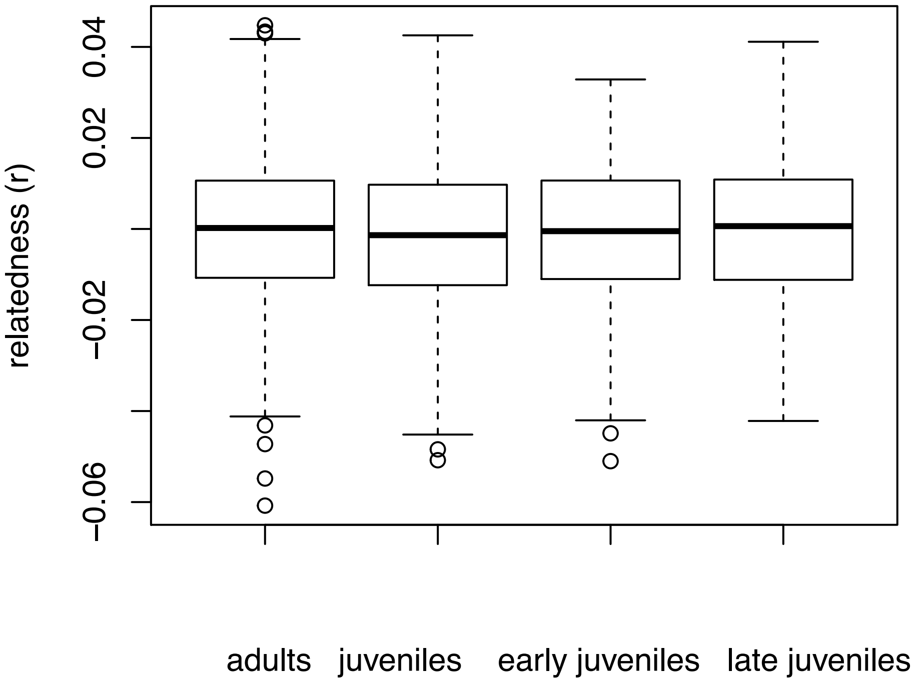


**Figure S5**: Box plots of relatedness values among different groups of individuals. Pairwise relatedness values do not decline across generations nor change within one recruitment season.

Figure S6:

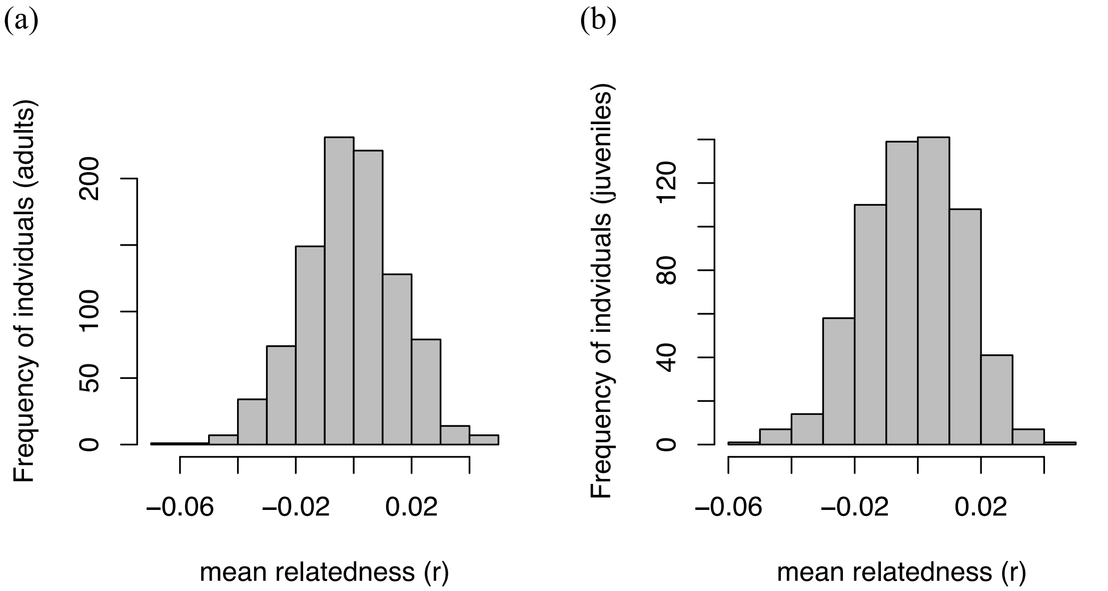


**Figure S6**: Frequency distribution of individual mean relatedness values for a) adults and b) juveniles. Note that general distriobution patterns of relatedness values do not differ across the two generations meaning no general increased or decreased genetic relationships over two years.

Figure S7:

a)

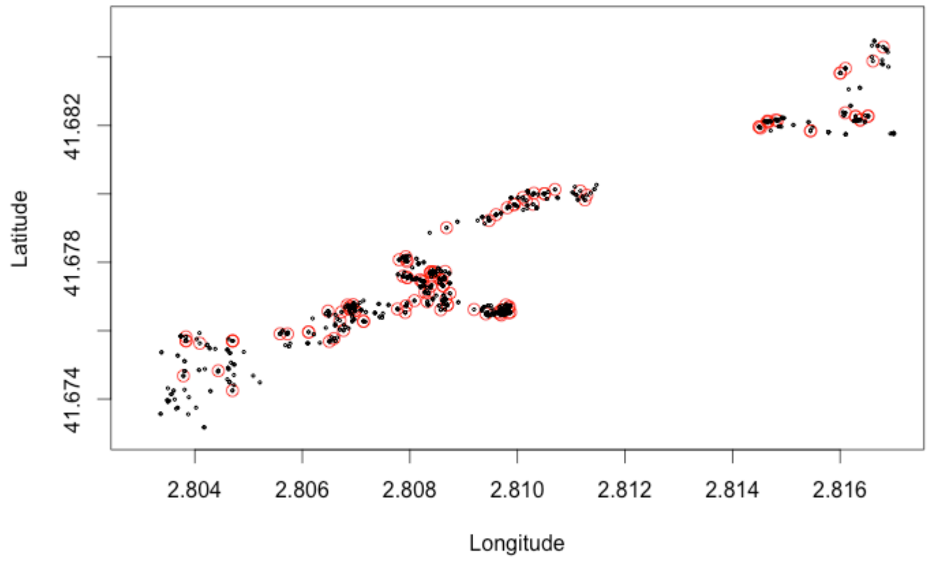


b)

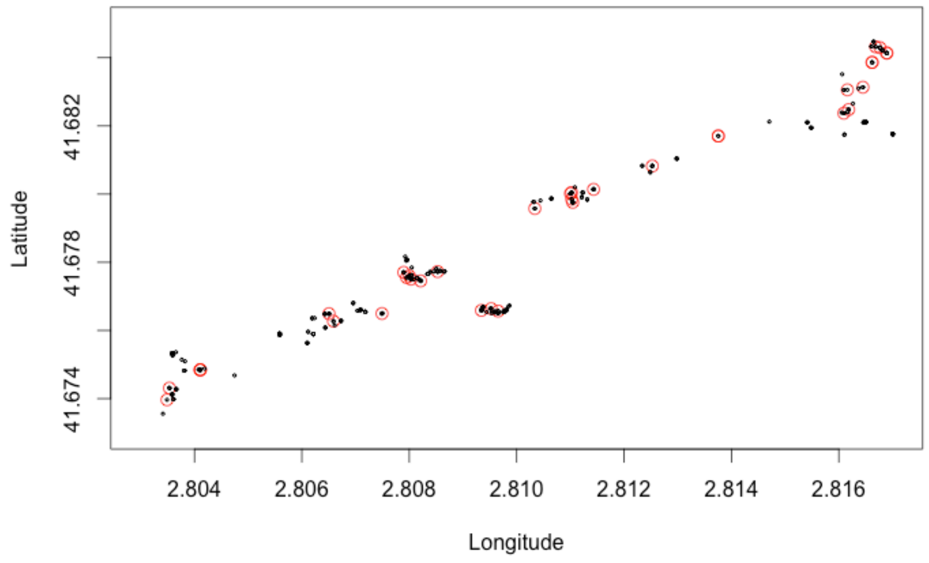


**Figure S7**: Spatial distribution of all individuals (black dots) and 'recent migrant' individuals (red dots) for adults (a) and juveniles (b) in the the small-scale area. 'Recent migrants' are the 25% of individuals (out of all adults or juveniles respectively) with the lowest ratios (bottom 25%). For adults it can be seen that there are less 'recent migrants' present in the southwestern part of the small sampling area.

Figure S8:

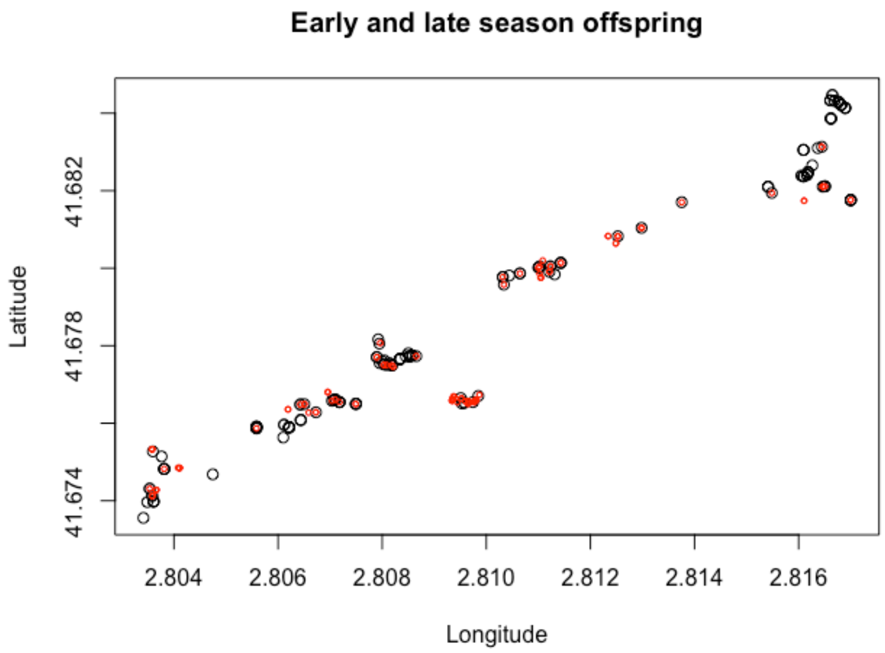


**Figure S8**: Spatial distribution of juveniles from the beginning of the season (black dots) and from the end of the season (red dots) in the small-scale area.

Table 1: Proportion of individuals in the small-scale area that belong to each of the four groups identified based on the combination of genetic differentiation within and between clusters (MIXED-range, SHORT-range dispersers, MEDIUM-range dispersers and HIGH-range dispersers) for each spatial cluster displayed in Figure 3

| **Adults** | SOUTH | | | | | | NORTH |
|---|---|---|---|---|---|---|---|
| | SplClust1 | SplClust2 | SplClust3 | SplClust4 | SplClust5 | | SplClust7 |
| N | 174 | 112 | 126 | 41 | 386 | | 107 |
| MIXED-range | 0 | 0.06 | 0 | 0 | 0 | | 0 |
| SHORT-range dispersers | **0.64** | **0.55** | **0.57** | **0.5** | 0.47 | | 0.39 |
| MEDIUM dispersers | 0.33 | 0.37 | 0.43 | 0.41 | **0.53** | | **0.54** |
| HIGH dispersers | 0.04 | 0.02 | 0 | 0.09 | 0 | | 0.07 |
| **Juveniles** | SplClust1 | SplClust2 | SplClust3 | SplClust4 | SplClust5 | SplClust6 | SplClust7 |
| N | 50 | 59 | 69 | 46 | 85 | 16 | 57 |
| MIXED-range | 0 | 0 | 0.2 | 0 | 0.26 | 0.5 | 0.12 |
| SHORT-range dispersers | 0.21 | 0.29 | 0.34 | 0.46 | **0.52** | 0.31 | **0.81** |
| MEDIUM-range dispersers | **0.79** | **0.71** | 0.36 | **0.54** | 0.12 | 0 | 0.07 |
| HIGH-range dispersers | 0 | 0 | 0.1 | 0 | 0.1 | 0.19 | 0 |
| **Early juveniles** | SplClust1 | SplClust2 | SplClust3 | SplClust4 | SplClust5 | SplClust6 | SplClust7 |
| N | 11 | 28 | 27 | 29 | 13 | 2 | 20 |
| MIXED-range | 0 | 0.11 | 0.29 | 0 | 0 | 0.26 | 0.15 |
| SHORT-range dispersers | 0.29 | 0.13 | 0.43 | 0 | **0.63** | 0.47 | 0.41 |
| MEDIUM-range dispersers | 0.41 | **0.53** | 0.29 | 0 | 0.38 | 0.26 | 0.36 |
| HIGH-range dispersers | 0.29 | 0.24 | 0 | 1 | 0 | 0 | 0.08 |
| **Late juveniles** | SplClust1 | SplClust2 | SplClust3 | SplClust4 | SplClust5 | SplClust6 | SplClust7 |
| N | 12 | 33 | 17 | 20 | 18 | 10 | 11 |
| MIXED-range | 0.30 | 0.41 | 0.3 | **0.58** | 0.28 | 0.5 | 0 |
| SHORT-range dispersers | 0.45 | 0 | 0 | 0.18 | 0.11 | 0 | **0.64** |
| MEDIUM-range dispersers | 0.25 | 0 | 0.4 | 0.12 | 0 | 0 | 0.36 |
| HIGH-range dispersers | 0 | **0.59** | 0.3 | 0.12 | 0.61 | 0.5 | 0 |

Clustering for adults only, juveniles only, early juveniles (those that settled early, before mid May, during the reproductive period) and late juveniles (those that settled late during the reproductive period, after mid June) is shown. SplClust = Spatial cluster corresponding to the clusters in Figure 3. Note that SplClust6 is not present in adults. MIXED-range dispersers: individuals with low Mean Genetic Distance (MGD) with respect to their own spatial cluster and low MGD with respect to alien spatial clusters. SHORT-range dispersers: individuals with low MGD with respect to their own spatial cluster and high MGD with respect to alien spatial clusters. MEDIUM-range dispersers: individuals with high MGD with respect to their own spatial cluster and low MGD with respect to alien spatial clusters. HIGH-range dispersers: individuals with high MGD with respect to their own spatial cluster and high MGD with respect to alien spatial clusters. Values above 0.5 are marked in bold.
